# Life stage associated remodeling of lipid metabolism regulation in the duplicated Atlantic salmon genome

**DOI:** 10.1101/140442

**Authors:** Gareth Gillard, Thomas N. Harvey, Arne Gjuvsland, Yang Jin, Magny Thomassen, Sigbjørn Lien, Michael Leaver, Jacob S. Torgersen, Torgeir R. Hvidsten, Jon Olav Vik, Simen R. Sandve

## Abstract

Atlantic salmon migrates from rivers to sea to feed, grow and develop gonads before returning to spawn in freshwater. These habitat shifts require great phenotypic plasticity. To address the unresolved question of how the shift in diet between fresh and saltwater affects the regulation of metabolic function, we fed salmon contrasting diets in each of the two life stages. Combining transcriptomics with comparative genomics, we found that lipid metabolism undergoes a concerted shift between fresh- and saltwater stages. Lipogenesis and lipid transport become less active in liver after transition to saltwater, while genes for lipid uptake in gut are more expressed in lipid-rich seawater environments. We assess how the whole-genome duplication that gave rise to the salmonids has impacted the evolution of lipid metabolism, and find signatures of pathway-specific selection pressure on gene duplicates, as well as a limited number of cases of increased gene dosage.

## Introduction

Atlantic salmon lives a ‘double life’. It starts its life in rivers, before transforming its physiology and behavior and migrating to sea to grow and accumulate resources for reproduction. This shift in environment requires preparatory remodeling of physiology prior to sea migration (referred to as smoltification), which encompasses a suite of coordinately regulated processes involving hormonal changes and large scale alteration of gene expression. The resulting adaptations to a marine environment include shifts in salt-tolerance, coloration, behavior, growth rate, and metabolism (reviewed in Stefansson et al., 2008).

Salmon transforms its lipid metabolism function during smoltification, likely related to differences in diet between environments (Sheridan, 1989). Salmon in rivers mostly eat invertebrates that are low in essential long-chain polyunsaturated fatty acids (LC-PUFA). Possibly as an adaptation to this (Leaver et al., 2008), salmon has evolved a greater capacity for endogenous production of LC-PUFAs than many other fish species, and several studies have demonstrated that salmon has the ability to increase or decrease endogenous omega-3 synthesis as a response to the dietary availability of LC-PUFAs (Kennedy et al., 2006; Leaver et al., 2008; Morais et al., 2011; Ruyter et al., 2000; Tocher et al., 2001; Tocher et al., 2002; Zheng et al., 2005). In contrast to rivers, marine habitat food chains are high in available LC-PUFAs, and it has therefore been hypothesized that smoltification-associated lipid metabolism changes are linked to preparation to this new dietary situation. However, little is known about the extent and nature of the lipid metabolism remodeling associated with the life stage shift from freshwater to sea.

The evolution of novel traits in salmonids, such as increased plasticity and the ability to migrate to sea, may have been facilitated by their ancestral whole genome duplication (called Ss4R) some 80 million years ago (Allendorf & Thorgaard, 1984; Lorgen et al., 2015; Macqueen & Johnston, 2014). Gene duplication can give rise to new adaptive phenotypes in different ways: through evolution of novel functions or gene regulation, subdivision and/or specialization of function among duplicates, or via an adaptive increase in gene dosage. The Atlantic salmon genome contains ~10,000 pairs of Ss4R gene duplicates, of which ~50% have evolved some novel regulation (Lien et al., 2016; Robertson et al., 2017). Indeed, in the context of lipid metabolism, it has recently been shown that a Ss4R duplicate of elovl5, a key enzyme in LC-PUFA syntheses, has gained expression compared to its ancestral regulation with likely implications for the ability to synthesize LC-PUFAs (Carmona-Antoñanzas et al., 2016). This is believed to have facilitated evolution of novel traits, including flexible phenotypes necessary for an anadromous life history (Stefansson et al., 2008). However, no systematic genome wide study has been conducted to assess the importance of the Ss4R in evolution of salmon lipid metabolism.

In this study, we integrate comparative genomics with transcriptomic data from feeding trials carried out across the fresh to saltwater transition to build a functional annotation of lipid metabolism pathway genes in salmon. We use this annotation to elucidate (i) the nature of the transformation of lipid metabolism from freshwater to saltwater life stages and (ii) the impact of whole genome duplication on evolution of the lipid gene repertoire and metabolic function. Our results indicate a programmed shift in lipid metabolism after transition to seawater, and show that lipid pathways differ with respect to selection pressure on gene duplicates from the salmonid whole genome duplication.

## Results and discussion

### Annotation of lipid metabolism genes

To identify genes involved in lipid metabolism in Atlantic salmon, we initially assembled groups of orthologous genes (orthogroups) from a selection of four salmonid species, Northern pike (their closest unduplicated relative), as well as teleost and mammalian outgroups (Figure 1a). Next, we aligned orthogroup proteins and constructed maximum likelihood gene trees. The majority (82-98%) of proteins from each species were represented in 23,782 ortholog gene trees. The salmonid species had significantly higher number of proteins included in ortholog gene trees compared to non-salmonid fish (Figure S1), reflecting the salmonid specific whole genome duplication. We then used the evolutionary distances in gene trees to infer the most likely salmon sequence orthologs of zebrafish genes selected from 19 KEGG pathways involved in lipid metabolism (File S1). This resulted in the annotation of 1421 (File S2) salmon lipid metabolism genes, of which 326 (23%) showed a 2:1 ortholog ratio between salmon and zebrafish (Figure 1b). Only 87 (6%) of the zebrafish genes could not be assigned a salmon ortholog.

**Figure 1:**
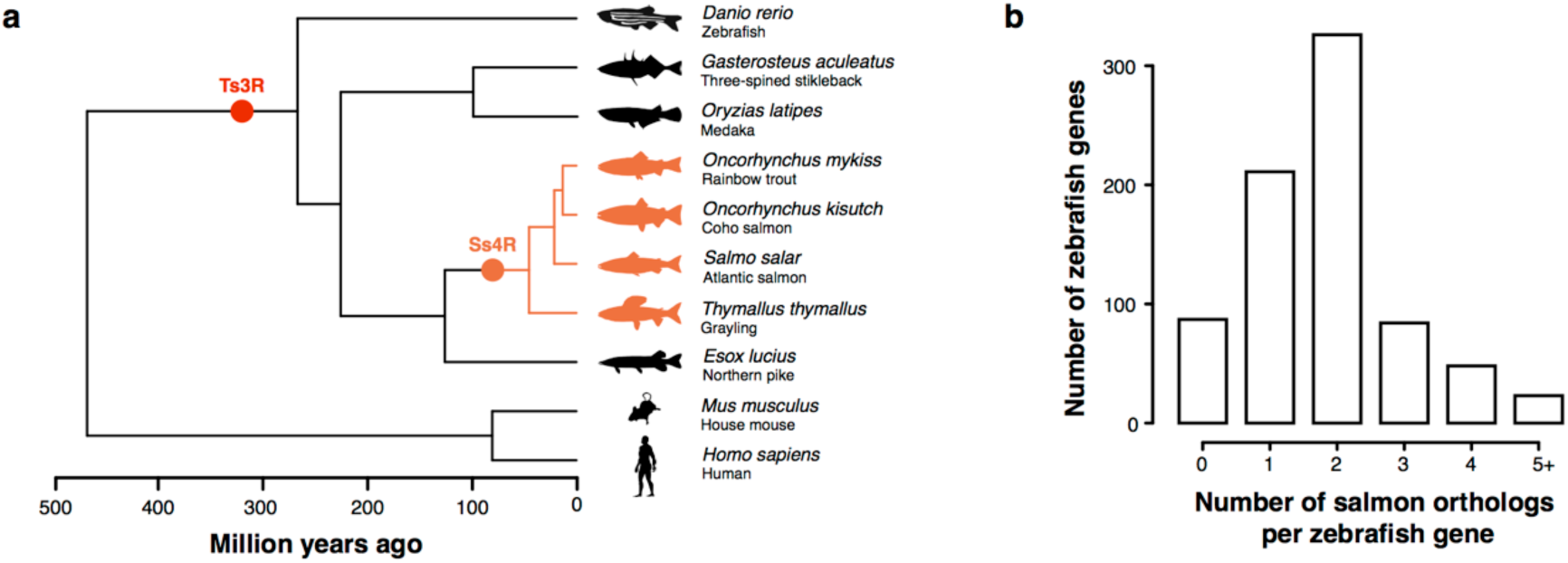
Ortholog annotation. (a) Species used to construct ortholog groups and their evolutionary distance. Points in the phylogenetic tree show the time of the teleost specific (Ts3R) and salmonid specific (Ss4R) whole genome duplications. (b) The number of salmon orthologs found (1421 genes in total) per zebrafish gene in 19 selected KEGG pathways involved in lipid metabolism.

To validate our ortholog annotation pipeline used to identify lipid metabolism genes, we analyzed the tissue specificity of these genes using gene expression data from 15 tissues in wild-type Atlantic salmon (File S3). Genes in certain fatty acid metabolism related pathways (*‘fatty acid metabolism’*, *‘PPAR signaling pathway’*, *‘fat digestion and absorption’*) had higher overall expression in tissues known to have high lipid metabolism activity (i.e. pyloric caeca, liver, and heart) (Glatz et al., 2010; Benedito-Palos & Pérez-Sánchez, 2016; Tocher, 2003) (Figure 2). Examples include: 1) Liver was the site of highest expression for all genes in the LC-PUFA biosynthesis pathway (the desaturases Δ6FAD and Δ5FAD, and the elongases elovl5, elovl2 and elovl4). 2) Bile acids are essential for fat digestion in the gut, but are synthesized in liver.

**Figure 2:**
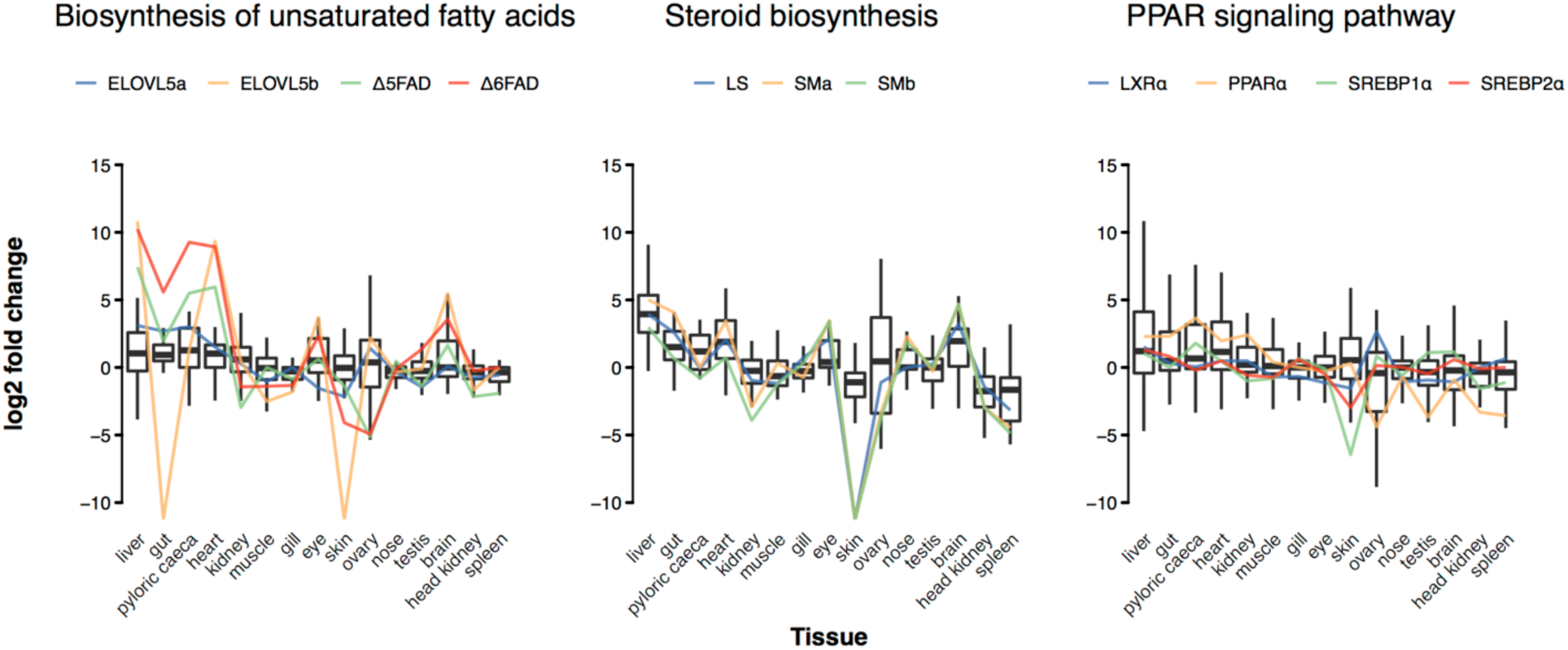
Tissue expression profiles of salmon genes in lipid metabolism pathways. Tissue expression profiles of our annotated lipid metabolism genes were consistent with expectations. Gene expression levels are shown as the log2 fold change difference between the FPKM value of each tissue and the median FPKM across all tissues. Expression profiles for selected genes in each pathway are shown (see Figure S2 and S3 for all pathways and gene details).

As expected, the rate limiting step for bile syntheses, cytochrome P450 7A1 (CYP7A1), has the highest expression in the liver. 3) Cholesterol, an essential component of cell membranes and precursor to bile acids, is known to be synthesized in all tissues, but primarily in liver, intestine, and brain (Brown & Sharpe, 2016). This is reflected in our annotation by high expression of the key cholesterol biosynthesis genes 3-hydroxy-3-methyl-glutaryl-CoA reductase (HMGCR), isopentenyl-diphosphase Aisomerase (IDI1), squalene epoxidase (SM), and lanosterol synthase (LS) in these tissues. 4) Several known regulators of lipid metabolism show high expression in liver, heart, brain and pyloric caeca, as expected, including liver X receptor (LXR), peroxisome proliferator-activated receptor alpha (PPARɑ), sterol regulatory element binding protein 1 (SREBP 1), and sterol regulatory element binding protein 2 (SREBP2). Taken together, the tissue distribution of lipid metabolism gene expression is in line with knowledge about vertebrate physiology in general, and support the validity of our annotation of lipid metabolism genes in salmon. To make all data underlying our annotation easily available, and to facilitate further refinement through manual community curation, we have created an interactive web-server available online (goo.gl/8Ap89a).

### Life-stage dependent remodeling of lipid metabolism

We conducted a feeding trial to study how salmon adjusts its lipid metabolism to different levels of LCPUFA in freshwater and saltwater. Groups of salmon were fed contrasting diets from hatching until after transition to seawater. One feed was vegetable oil based (VO) and hence low in LC-PUFA, similar to river ecosystem diets, whereas the other was based on fish oil (FO) and high in LC-PUFA as expected in a marine-type diet (see Table S3 for details on feed composition). VO based diets are also known to contain lower cholesterol levels (Ciftci, et al., 2012; Verleyen et al., 2002). In total, 32 and 23 fish were sampled for RNA-seq of liver and gut, respectively, including eight biological replicates from each diet and life-stage (freshwater and saltwater). Fish in the different dietary groups were given FO and VO feed from first feeding (<0.2g) until sampling.

In general, global gene expression levels were more affected by dietary composition in liver than in gut (which was largely unresponsive), and the effect was more pronounced in freshwater than in saltwater (Figure 3a). VO diets increased lipid-metabolism related gene expression compared to FO diets, with 57 genes (86% of the differentially expressed genes, DEGs) upregulated in freshwater and 23 genes (74% of DEGs) upregulated in saltwater (Figure 3b). The increased activity of liver lipid metabolism under VO diets confirm the well-known ability of salmon to regulate endogenous synthesis of LC-PUFA and cholesterol in response to VO diets (Kortner et al., 2014; Leaver et al., 2008; Zheng et al., 2005).

**Figure 3:**
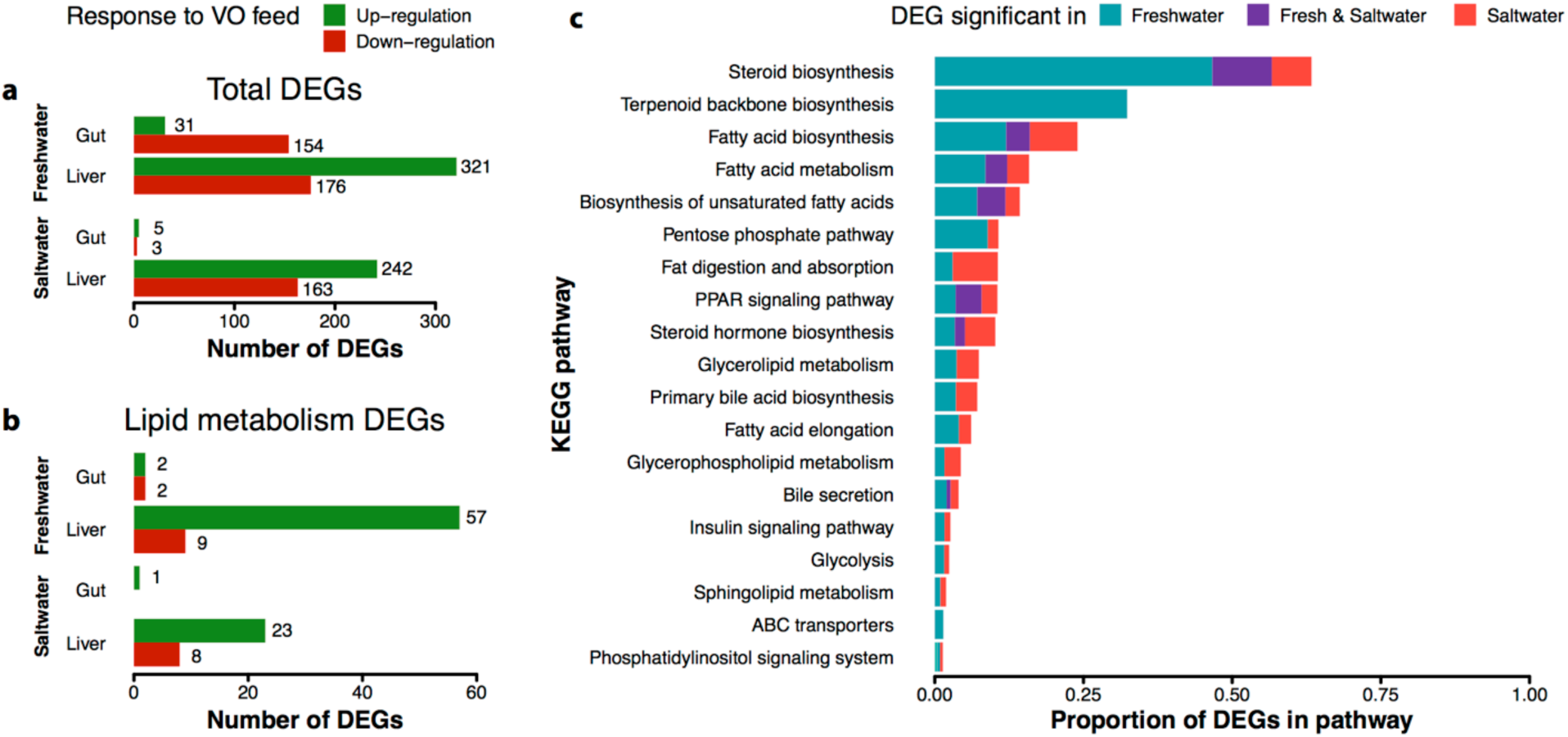
Gene regulation in response to feed type. (a) Total number of significant (FDR < 0.05) differentially expressed genes (DEGs) between fish oil (FO) and vegetable oil (VO) fed salmon in the liver and gut tissues of freshwater and saltwater stage Atlantic salmon (see Files S4 (liver) and S5 (gut) for underlying data). (b) As above, but for lipid-associated genes only. (c) Proportions of genes in each KEGG pathway that had significantly different liver expression between the two feed types only in freshwater, only in saltwater, or in both stages.

Fish sampled in freshwater and saltwater shared a relatively small number of DEGs for each pathway (Table S4). We found that some pathways had more DEGs in freshwater (‘*fatty acid biosynthesis’*, ‘*steroid biosynthesis*’, and its precursor ‘*terpenoid backbone biosynthesis’*), whereas others had more DEGs in saltwater (‘*steroid biosynthesis’*, ‘*fatty acid biosynthesis’*, and ‘*fat digestion and absorption’*) (Figure 3c). Out of 71 lipid metabolism DEGs in the dietary contrast, 78% (51 genes) were freshwater specific, 11% (8 genes) saltwater specific, and 11% (8 genes) shared dietary response (Table S4). For example, only two genes in the FA and LC-PUFA biosynthesis pathways (Δ6FADa and Δ5FAD) shared response to dietary availability in fresh- and saltwater (Figure 4). A similar trend was found for diet-induced expression changes in the pathways responsible for cholesterol biosynthesis (Figure 5). The few genes that showed diet-effects specific to saltwater included bile salt activated lipase, responsible for the hydrolysis of free fatty acids from TAG obtained from the diet (Tocher, 2003). Two of these genes, carboxyl ester lipase, tandem duplicate 2a (CEL2a) and b (CEL2b), are highly upregulated in saltwater in response to VO diet. Taken together, our results show clear evidence of both life-stage- and diet-dependent remodeling of endogenous liver lipid metabolism and corroborates the idea of a post-smoltification phenotype adapted to an environment with a surplus of essential lipids.

**Figure 4:**
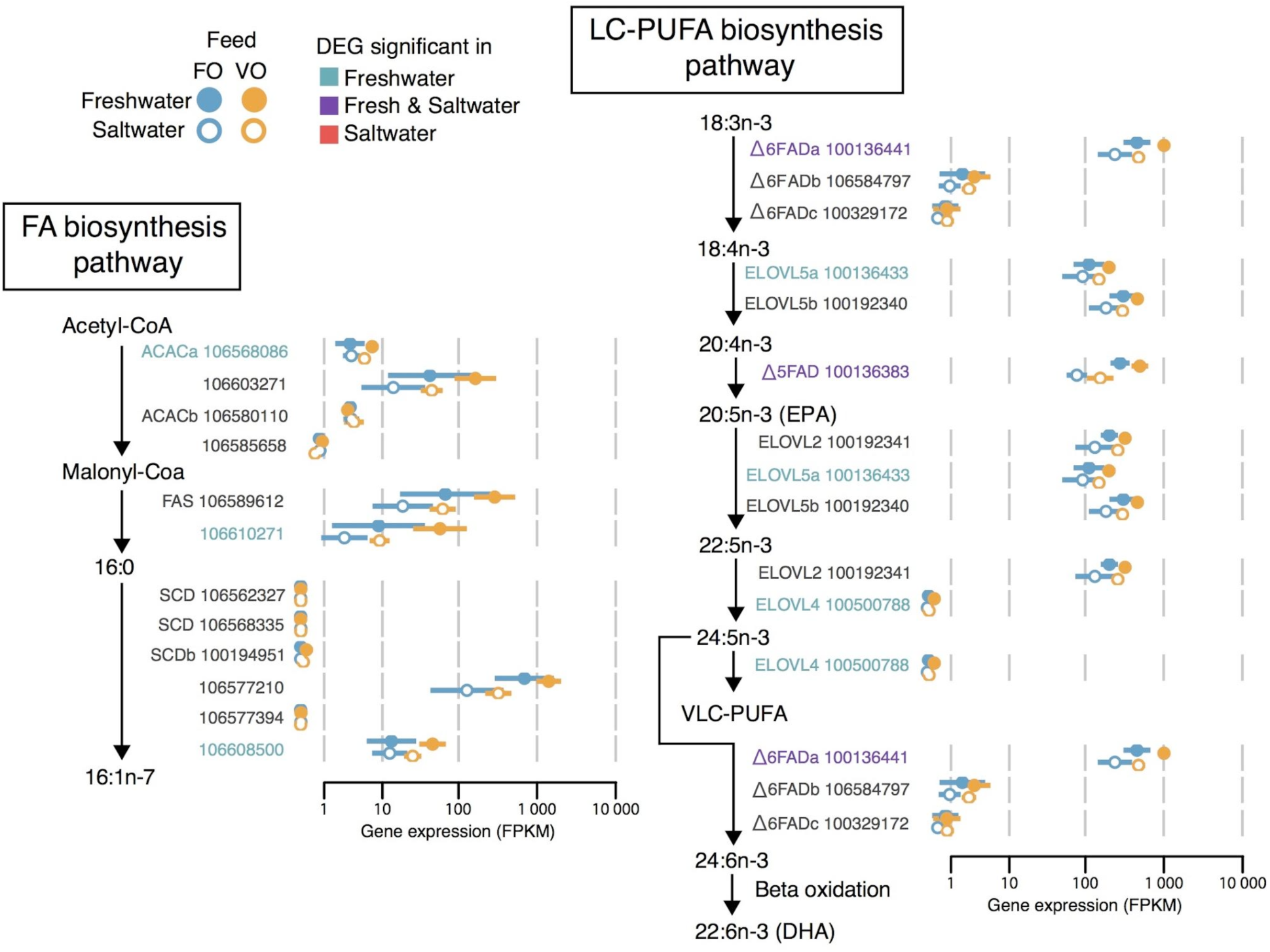
Diet and life stage effects on FA and LC-PUFA biosynthesis in salmon liver. Core fatty acid (FA) biosynthesis and biosynthesis of unsaturated fatty acids pathways with Atlantic salmon genes annotated to each catalytic step (enzyme names followed by NCBI gene numbers). Gene expression levels are shown as mean (point) and standard deviation (line) of expression in eight samples (measured in log(FPKM + 1)) from each diet (FO, VO feeds) and life stage (freshwater, saltwater) combination. Genes significantly (FDR<0.05) differentially expressed (DEG) between diets in a life stage are highlighted.

**Figure 5:**
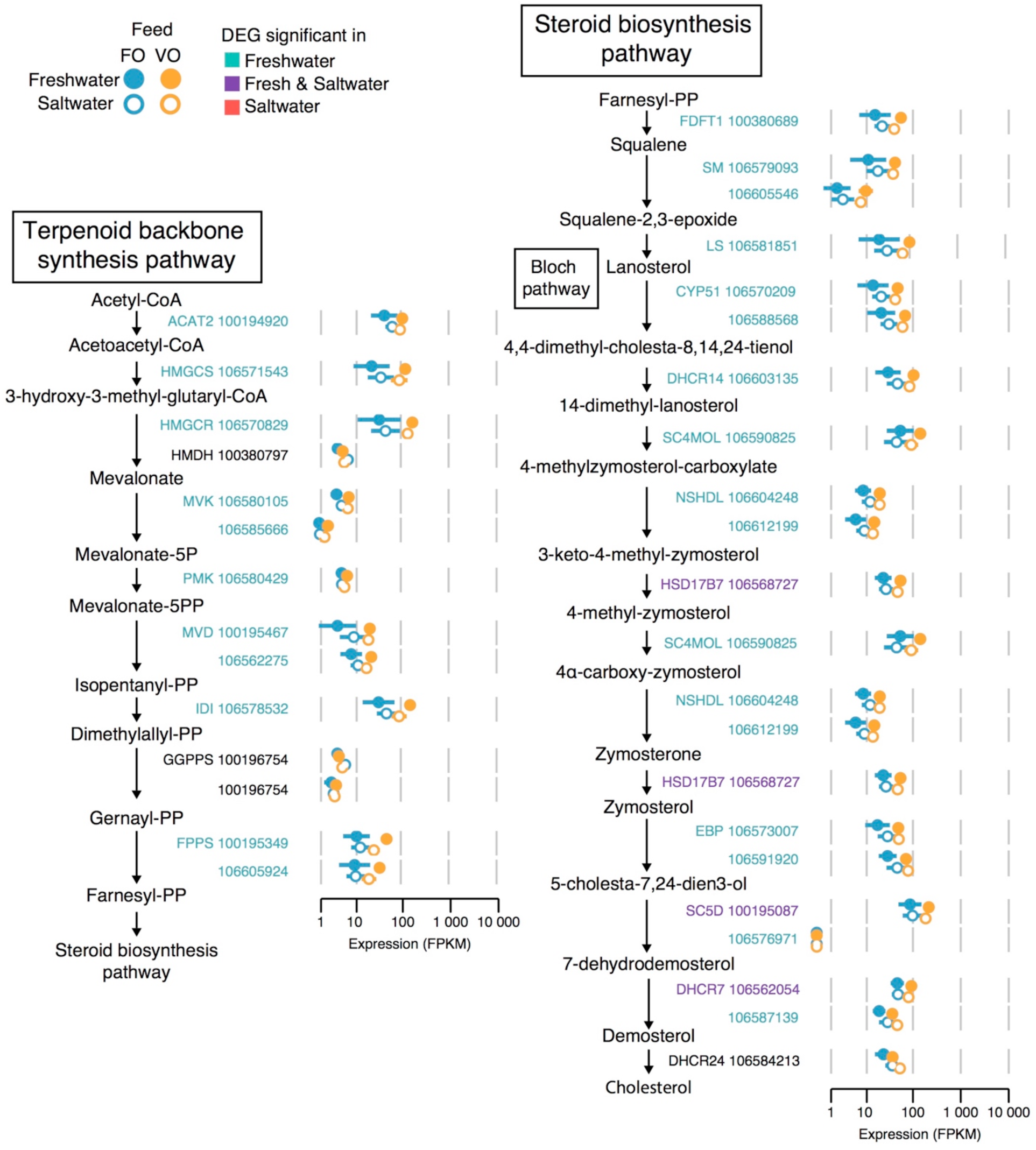
Diet and life stage effects on cholesterol biosynthesis in salmon liver. Terpenoid backbone synthesis and steroid biosynthesis pathways with Atlantic salmon genes annotated to each catalytic step (enzyme names followed by NCBI gene numbers). Gene expression levels are shown as mean (point) and standard deviation (line) of expression in eight samples (measured in log(FPKM + 1)) from each diet (FO, VO feeds) and life stage (freshwater, saltwater) combination. Genes significantly (FDR<0.05) differentially expressed (DEG) between diets in a life stage are highlighted.

To further investigate the life-stage associated changes in lipid metabolism we tested for differential expression between freshwater and saltwater in salmon fed the same diets (Figure 6). Liver and gut showed contrasting effects of saltwater on lipid gene expression with extensive downregulation in liver and upregulation in gut (Figure 6b). The number of DEGs in each tissue were similar for the environment comparison (Figure 6a), unlike for the diet comparison (Figure 3).

**Figure 6:**
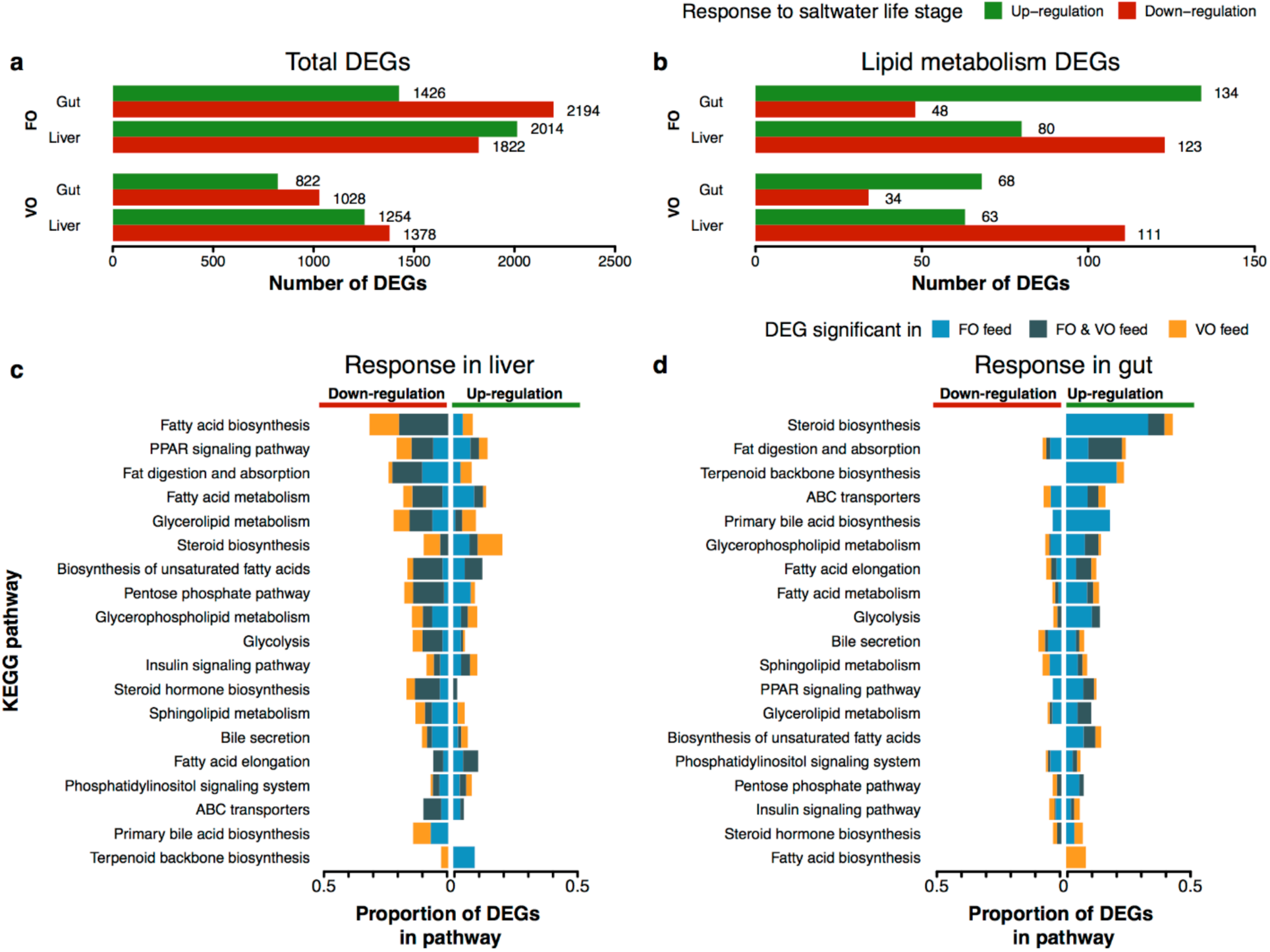
Gene regulation in response to life stage. (a) Total number of significant (FDR < 0.05) differentially expressed genes (DEGs) between freshwater and saltwater life stages in the liver and gut tissues of Atlantic salmon fed fish oil (FO) or vegetable oil (VO) diets (see Files S6 and S7 for underlying data). (b) As above, but for lipid metabolism DEGs. (c) Proportion of genes in each KEGG pathway that are DEGs in liver and (d) gut, colored by DEG significance in only FO, only VO, or both diets, and separated into up- or down-regulation in saltwater samples.

Further examination of key lipid metabolism genes revealed that during smoltification the system-wide lipid metabolism remodeling represented a concerted shift in the metabolic role of liver and gut. As the salmon entered the marine stage, lipogenic gene expression in the liver was significantly decreased, as evident by the markedly lower expression (2.2-3.3 fold) of the master regulator of lipid metabolism SREBP1, a 5-fold decrease in expression of fatty acid synthase, and a 2-3 fold decrease in rate-limiting enzymes in LC-PUFA synthesis (i.e. Δ5FAD, Δ6FADa) (Figure 4). Liver and gut gene expression also indicated increased catabolic activity in saltwater, upregulating the uptake of fatty acids into mitochondria for β-oxidation (uptake carried out by carnitine palmitoyltransferase 1 and 2) (Lehner & Quiroga, 2016). Finally, expression of lipid transport genes shifted from liver to gut with the transition to seawater (apolipoproteins, pathway “Fat digestion and absorption” in Figure 6). Four apolipoproteins (out of 11 annotated) were differentially regulated in liver between different, with a 2.4-5 fold decrease in saltwater compared to freshwater. In stark contrast, nine of the diet-regulated apolipoproteins in gut increased their expression in saltwater between 1.8-9.7 fold. These results suggest that the decreased ability of Atlantic salmon to synthesize LC-PUFAs after fresh-to-saltwater transition is compensated by an increased ability to take up lipids in the gut.

Interestingly, diet had a strong influence on the number and direction of gene expression changes between freshwater and saltwater (Figure 6). In gut, the transcriptional changes between fresh- and seawater was twice as high when fed FO diet than VO diet (Figure 6a). In liver, the diet effect was less pronounced, with the FO group containing 46% more DEGs than the VO group (Figure 6a). Identical patterns were found for lipid metabolism genes with 89% and 16% more DEGs in FO group for gut and liver, respectively (Figure 6b). As this diet and life-stage interaction is a genome wide trend, and more pronounced in gut tissue than in liver, this pattern could be related to differences in osmoregulation and adaptation to saltwater. The higher levels of essential omega-6 FA linoleic acid (18:2 n-6, LA) in VO diets could result in increased levels of arachidonic acid (20:4 n-6, ARA). ARA is a precursor to eicosanoids with a multitude of biological functions including osmoregulation, a critical function of gut (Mustafa & Srivastava, 1989; Oxley et al., 2010). Another possibility is that the different levels of fatty acids in the diets, for example DHA, affect DNA-methylation and thus trigger genome wide divergence in gene regulation (Kulkarni et al., 2011).

Our results clearly demonstrate very different baseline lipid metabolic functions in pre- and post-smolt salmon, as well as life-stage associated changes in the plasticity of lipid metabolism, e.g. the ability to regulate endogenous LC-PUFA synthesis as a response to changes in diet (i.e. lipid content). As opportunistic carnivores, salmon tend to eat whatever the local environment provides. Thus, in freshwater, insects and amphipods provide variable, mostly low amounts of essential LC-PUFA (Jonsson & Jonsson, 2011; Sushchik et al., 2003), favoring a metabolic function that can efficiently regulate endogenous lipid synthesis based on dietary availability (Carmona-Antonanzas et al., 2014). Conversely, marine environments provide a stable source of essential dietary lipids, promoting a metabolic function that allocate less energy to endogenous synthesis of essential lipids.

### Selection on gene duplicates after whole genome duplication

Carmona-Antonanzas et al. (2014, 2016) proposed that the salmonid whole-genome duplication may have adaptively increased the potential for endogenous lipid synthesis. We pursued this hypothesis by searching for distinct signatures of selection pressure on lipid metabolism genes in salmon. Specifically, we compared pathways in terms of their tendency to retain both duplicates of gene pairs, in terms of whether duplicates showed similar regulation (expression patterns across tissues, diets and environments), and in terms of total gene dosage (for the one or two genes retained of a pair) in salmon compared to pike, its closest unduplicated sister lineage.

To assess the level of Ss4R duplicate retention, we first defined 10,752 Ss4R duplicate pairs (21,504 genes) in the NCBI refseq annotation using the same approach as Lien et al. (2016). Of the 1,421 annotated lipid metabolism genes, 867 (61%) were retained as duplicated genes after Ss4R (Figure 7a) (in contrast to 47% of the 45,127 salmon genes assigned to ortholog groups). Moreover, our results showed large variation in the proportion of retained duplicates in each lipid metabolism pathway (Figure 7), with the most extreme case being ‘*fat digestion and absorption*’ with 80% retained duplicates and ‘*steroid hormone biosynthesis*’ with only 27% retained Ss4R duplicates.

**Figure 7.**
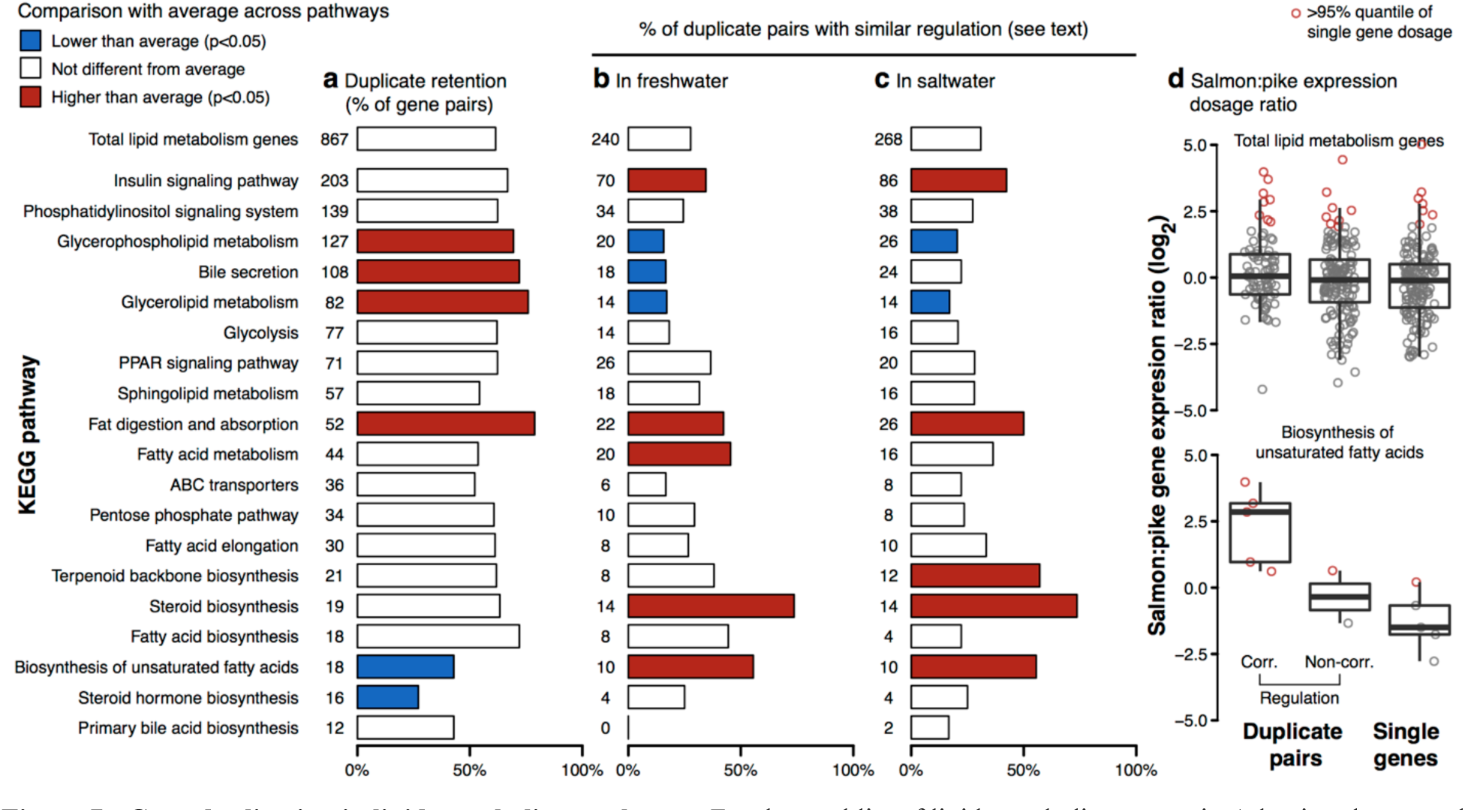
Gene duplication in lipid metabolism pathways. For the total list of lipid metabolism genes in Atlantic salmon, and sets of genes belonging to different KEGG pathways: (a) Number and percentage of genes with a duplicate homolog from the Ss4R duplication. (b) Number and percentage of duplicate genes with correlated liver expression response to feed in freshwater and (c) saltwater (Correlation >= 0.6, P-value < 3.306e-3, using 19 time points from feed trial for each water condition). Fisher’s exact test was used to detect pathways with significant enrichment compared to all gene (P-value < 0.05) (d) Log2 gene dosage ratios (salmon:pike) in liver from fish in freshwater, where the ratio is computed between expression in the salmon duplicates (FPKM, sum of the two duplicates) and the expression of the corresponding pike ortholog. Ratios were computed for all lipid metabolism genes and genes in the pathway *‘biosynthesis of unsaturated fatty acids’*. For comparison, ratios were also computed for genes without retained duplicates, i.e. with a 1:1 orthology between salmon and pike. Duplicates were grouped into correlated (corr.) or non-correlated (non-corr.) based on saltwater correlation result in (c). Dosage ratios (points) greater than the 95% quantile of single gene dosages are marked in red.

The regulatory conservation of the duplicates was then estimated from RNA-seq data representing a time course of dynamic changes in gene expression and lipid metabolism function in liver. Fish in the same feeding trial (seen description above) were switched from VO to FO feed and vice versa, in both fresh and saltwater conditions (see methods and supplementary data for details). In total, 38 sampling time points (19 in freshwater and 19 in saltwater) from the switch experiment were used to calculate co-expression correlation for Ss4R duplicates. Pathway-level analyses showed that conservation at the regulatory level was not associated with duplicate retention (Figure 7). For example, the *‘biosynthesis of unsaturated fatty acids’* pathway had significantly fewer duplicates retained than expected by chance (P-value < 0.0234), but a significant overrepresentation of duplicate pairs that display highly similar regulation (P-value < 0.0142 and < 0.0361 in freshwater and saltwater, respectively). Other pathways showing signatures of increased duplicate co-regulation were *‘insulin signalling pathway’*, *‘terpenoid backbone biosynthesis’*, *‘steroid biosynthesis’*, *‘fat digestion and absorption’*, and *‘fatty acid metabolism’* (Figure 7b-c). Overall, the distinct differences in duplicate retention and conservation of regulatory mechanisms across the lipid metabolism pathways suggest differences in selective pressures shaping duplicate evolution following Ss4R. Moreover, the pathways with highly conserved duplicate co-regulation were also those that were most responsive to dietary differences in fatty acid profile (Figure 3).

Finally, to link duplicate retention and co-regulation to signals of increased gene dosage following Ss4R, we used RNA-seq data from the Northern pike (*Esox lucius*), a species that belongs to the unduplicated sister lineage (see methods for details). For each duplicate pair, we computed the ratio between the sum of Ss4R duplicate expression and its non-duplicated ortholog in pike and compared these ratios to those observed for salmon genes that had not retained two Ss4R duplicates. In total 69 duplicate pairs from 18 different lipid-metabolism related pathways displayed a combined dosage increase relative to single copy genes, of which 26 had highly conserved regulation (i.e. correlated expression) (File S8). We saw no systematic effect of gene dosage when comparing the total gene expression of duplicate pairs with that of single-copy genes; nor did co-regulation of duplicates associate with increased gene dosage (Figure 7d). This pattern was also true for most individual lipid pathways (Figure S4-S5), except for *‘biosynthesis of unsaturated fatty acids’*, *‘fatty acid metabolism’* and *‘fatty acid elongation’*. These three pathways showed a link between co-regulation of duplicated genes and higher total gene dosage (Figure S4-S5, Figure 7d). Underlying this link were three genes with co-regulated dosage effects shared between all three pathways; trifunctional enzyme alpha subunit b (hadhab), elovl6, and the previously identified elovl5 (Carmona-Antonanzas et al., 2014; Carmona-Antoñanzas et al., 2016). Only elovl5 is known to be directly involved in core PUFA biosynthesis. Hadhab is involved in mitochondrial β-oxidation/elongation and elovl6 is involved in elongation of saturated and monounsaturated fatty acids (Bond et al., 2016). Although we do not see a general trend of increased gene dosage effects on lipid metabolism genes after whole genome duplication, it is likely that an increased dosage of *elovl5* and the 68 other duplicate pairs has affected the function of lipid metabolism in salmon.

## Conclusion

Atlantic salmon needs great plasticity of physiology and behavior to adapt for migration between freshwater and sea. By analyzing transcriptomic changes through the transition from fresh- to saltwater, we identified an overall remodeling of lipid metabolism, with liver more active in freshwater and gut more active in saltwater. This baseline remodeling was modulated by diet, as life-long dietary contrasts showed that lipid metabolism was significantly more responsive to dietary differences in lipid composition in fresh water versus saltwater. These results indicate adaptive optimization of the Atlantic salmon lipid metabolism to account for life-stage specific dietary availability. Moreover, we found signatures of pathway-specific selection pressure on gene duplicates, including a gene dosage increase in three genes involved in fatty acid metabolism. This illustrates possible adaptive consequences of the salmonid whole-genome duplication for the evolution of lipid metabolism. Future studies should attempt to decipher how this life-stage related metabolic reprogramming is controlled. Elucidating the extent and mechanisms to which physiological transformation before sea migration in salmonids is “hard coded” (for example through epigenetic remodeling) will further understanding of evolutionary processes, and has economically important implications for aquaculture.

## Materials and methods

### Orthogroup prediction

Protein sequences were obtained from seven teleost fish species; *Danio rerio* (zebrafish), *Gasterosteus aculeatu* (three-spined stickleback), *oryzias latipes* (medaka), *Oncorhynchus mykiss* (Rainbow trout), *Oncorhynchus kisutch* (coho salmon), *Salmo salar* (Atlantic salmon), *Thymallus thymallus* (grayling), *Esox lucius* (northern pike), and two mammalian outgroup species; *Homo sapiens* (human), *Mus musculus* (house mouse). Human, mouse, zebrafish, medaka and stickleback protein fasta data were obtained from ENSEMBL (release 83). Atlantic salmon (RefSeq assembly GCF_000233375.1, Annotation Release 100) and northern pike (RefSeq assembly GCF_000721915.2, Annotation Release 101) proteins were obtained from NCBI RefSeq. Rainbow trout proteins were obtained from an assembly and annotation of the genome (Berthelot et al., 2014). Grayling proteins were obtained from an assembly and annotation of the genome (Varadharajan *et al.*, awaiting publication of data on NCBI/biorxiv in end of June 2017). The coho salmon transcriptome (Kim, Leong, Koop, & Devlin, 2016) was obtained from NCBI (GDQG00000000.1). Where transcriptome data was used, protein sequences were translated using TransDecoder (v2.0.1, http://transdecoder.github.io/). Protein fasta files were filtered to retrieve only the longest protein isoform per gene. Orthofinder (v0.2.8) (Emms et al., 2015) assigned groups of orthologs based on protein sequence similarity. Proteins within an orthogroups were further aligned using MAFFT (v7.130) (Katoh et al., 2002) and maximum likelihood trees were estimated using FastTree (v2.1.8) (Price et al., 2010).

### Annotation of salmon lipid metabolism genes

A list of zebrafish proteins obtained from 19 manually selected zebrafish KEGG pathways related to lipid metabolism (Table S1) were used to search for Atlantic salmon orthologs. Orthogroups that contained a selected zebrafish protein were identified. Salmon proteins within those orthogroups were assigned as orthologs of the closest zebrafish protein based on the orthogroup tree distance. A lipid metabolism gene list was created including salmon orthologs to the selected zebrafish genes. Additional salmon genes related to lipid metabolism not included in KEGG pathways (e.g. regulators or transporters, SREBP, LXR, FABP, etc.) were manually searched for through NCBI and added to the list.

### Tissue expression

Atlantic salmon RNA-seq samples from 15 different tissues (liver, gut, pyloric caeca, heart, kidney, muscle, gill, eye, skin, ovary, nose, testis, brain, head kidney, spleen) were obtained from NCBI SRA (PRJNA72713) (Lien et al., 2016). Fastq files were adapter trimmed before alignment to the Atlantic salmon genome (RefSeq assembly GCF_000233375.1) (Lien et al., 2016) using STAR (v2.5.2a) (Dobin et al., 2013). HTSeq-count (v0.6.1p1) (Anders et al., 2015) counted the sum of uniquely aligned reads in exon regions of each gene in the annotation (RefSeq Annotation Release 100). Gene FPKM values were calculated based on the gene count over the samples effective library size (see TMM method from edgeR (Robinson et al., 2010) user manual) and the mean gene transcript isoform length.

### Feed trial

Atlantic salmon fry were obtained from AquaGen Breeding Centre, Kyrksæterøra, Norway and reared in the Norwegian Institute for Water Research (NIVA), Solbergstranda, Norway on vegetable oil (VO) or fish oil (FO) based diets from first feeding (fry weight <0.2 g). VO based feeds contained a combination of linseed oil and palm oil at a ratio of 1.8:1 and FO based feeds contained only North Atlantic fish oil. All feeds were formulated and produced by EWOS innovation (Supplementary File 3). Smoltification was triggered by 5 weeks of winter-like conditions with 12 hours of light per day followed by spring-like conditions with 24 hours of light per day. Salmon were then immediately switched to saltwater and allowed to acclimate for 3 weeks before first sampling. Pre-smolt salmon in freshwater (~50 g) and post-smolt salmon in saltwater (~200 g) were either switched to the contrasting feed condition (VO to FO and vice versa) or maintained on the same feed as a control. Fish samples were taken on days 0, 1, 2, 6, 9, 16, and 20 post each diet switching, with five fish from each of two replicate tanks per treatment (5*2). Feeding was stopped in the mornings of each of the sampling days. All fish were euthanized by a blow to the head and samples of liver and midgut (gut section between pyloric caeca and hindgut) were flash frozen in liquid nitrogen and stored under -80 °C for further analysis.

### RNA-sequencing

Total RNA was extracted from selected feed trial samples using the RNeasy Plus Universal kit (QIAGEN). Quality was determined on a 2100 Bioanalyzer using the RNA 6000 nano kit (Agilent). Concentration was determined using a Nanodrop 8000 spectrophotometer (Thermo Scientific). cDNA libraries were prepared using the TruSeq Stranded mRNA HT Sample Prep Kit (Illumina). Library mean length was determined by running on a 2100 Bioanalyzer using the DNA 1000 kit (Agilent) and library concentration was determined with the Qbit BR kit (Thermo Scientific). Single end sequencing of sample libraries was completed on an Illumina HiSeq 2500 with 100 bp reads.

### Differential expression analysis of feed and life stages

To analyze gene expression differences between feed types and life stages, samples from the feed trial were selected for RNA-seq. The liver and gut tissue of a number of replicate fish were sequenced for each of the feeds (FO, VO) at day 0 of the diet switch, both before (freshwater) and after (saltwater) smoltification. Fastq files were processed to produce gene count and FPKM data using the same protocol described under the tissue expression method section. For feed comparison, changes in gene expression were tested between FO and VO feeds for both freshwater and saltwater samples, and liver and gut tissues. For life stage (water condition) comparison, changes in gene expression were tested between freshwater and saltwater stages for both FO and VO feed samples, and liver and gut tissues. Using RNA-seq gene count data, lowly expressed genes were filtered prior to testing, retaining genes with a minimum of one read count per million (CPM) in two or more samples. Differential expression analysis was carried out using a standard edgeR (Robinson et al., 2010) protocol. Effective library sizes were calculated using the edgeR TMM-normalisation procedure allowing effective comparison of expression data between different sample types (see edgeR user manual). An exact test between expression levels of a pair of conditions gave the log2 fold change, P-value and false discovery rate (FDR) for each gene. Genes with an FDR < 0.05 were considered a differentially expression gene (DEG).

### Identification of Ss4R duplicates

To identify putative gene duplicates stemming from the Ss4R we used the same approach as in Lien et al. (2016). All-vs-all protein blast was run with e-value cutoff of 1e-10 and pident (percentage of identical matches) ≥80 and blast hit coverage of ≥50% of protein length. Only the best protein hits between the 98 defined synteny blocks (see Lien et al., 2016) were considered as putative Ss4R duplicates. Blast result ranking was done using the product of pident times bitscore to avoid spurious ‘best blast matches’ with low pident (<85) but high bitscore.

### Duplicate analysis

Genes from the lipid metabolism gene list were paired together with their putative Ss4R duplicates identified above. The retention of gene duplicates (i.e. whether both genes in a pair were retained, or just one) was compared between all identified duplicates in the salmon genome annotation and the lipid metabolism gene list. Pathway-level retention was explored by comparing the number of genes in each of the 19 selected KEGG pathways (Table S1) in a duplicate pairing to that of the total list of lipid genes, to find pathways with significantly less or more duplicate retention (Fisher’s exact test, P-value < 0.05). Regulatory conservation of lipid gene duplicates was explored by correlation of gene expression changes between duplicates over the course of the feed trial described above. RNA-seq data was generated from liver samples of salmon from 38 sampling time points (19 in freshwater and 19 in saltwater). Fastq files were processed to produce gene count and FPKM data using the same protocol described under the tissue expression method section. For each duplicate pair, mean FPKM values were retrieved for each time point and used to calculate a freshwater and saltwater correlation value. Duplicates with Pearson correlation ≥ 0.6 were considered correlated (P-value < 0.003 from 19 sample points). The number of duplicates with correlated expression profiles was counted for each pathway and compared to all lipid genes to find pathways with significantly less or more correlated duplicates (Fisher’s exact test, P-value < 0.05). The effect of gene duplication on gene dosage was estimated by calculating a dosage ratio between the FPKM value of a salmon ortholog (sum of gene expression in duplicate pairs) over the FPKM value of the non-duplicated ortholog from northern pike. For salmon, the RNA-seq data from the freshwater and saltwater FO fed trial was used (samples used in differential expression analysis section). For pike, RNA-seq from livers of four individuals were aligned (see tissue expression section for protocol) to their respective genomes (see genomes in ortholog prediction section). RSEM (v1.2.31) (Li & Dewey, 2011) was used to generate FPKM values for genes so that non-uniquely mapped reads between salmon duplicate genes were not ignored but instead assigned proportionately to each gene to match the proportions of uniquely mapped reads between the genes. Gene dosage levels for duplicate pairs with correlated expression (see above), non-correlated expression and single genes was compared for all lipid metabolism genes and for each pathway.

## Supplemental material

**Figure S1:**
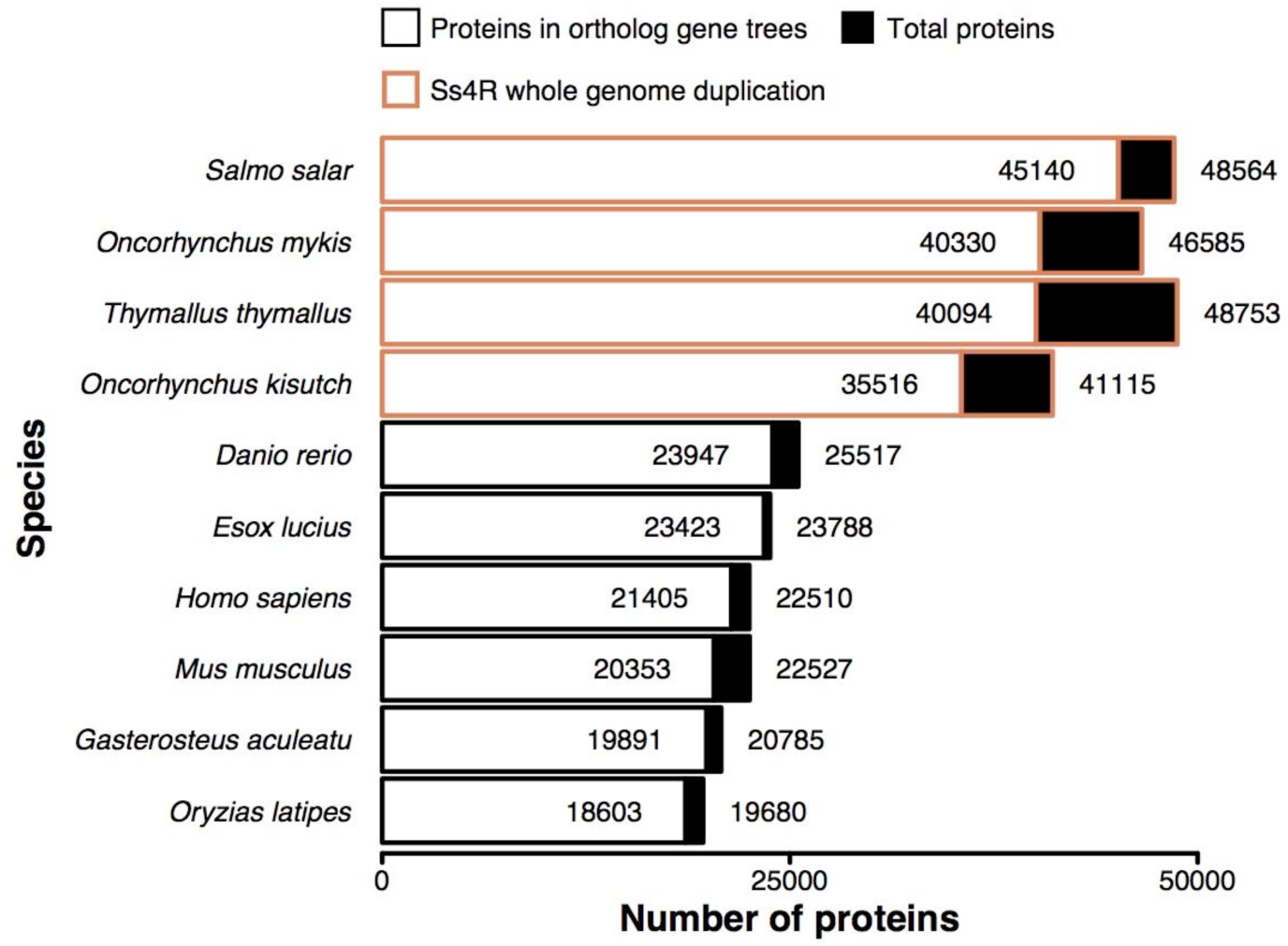
Number of proteins in ortholog gene trees for each species.

**Table S1:** KEGG pathways related to lipid metabolism selected for gene annotation. Identifiers prefixed with “dre” refer to *Danio rerio* (zebrafish), while “ko” (KEGG orthology) are generic (species-independent). The salmon pathways (prefix “sasa”) were not available when this paper was written.

| KEGG ID | Pathway name |
| --- | --- |
| dre00010 | Glycolysis |
| dre00561 | Glycerolipid metabolism |
| dre04070 | Phosphatidylinositol signaling system |
| dre04910 | Insulin signaling pathway |
| dre00600 | Sphingolipid metabolism |
| dre01212 | Fatty acid metabolism |
| dre03320 | PPAR signaling pathway |
| dre00564 | Glycerophospholipid metabolism |
| ko04976 | Bile secretion |
| dre02010 | ABC transporters |
| dre00900 | Terpenoid backbone biosynthesis |
| ko04975 | Fat digestion and absorption |
| dre00100 | Steroid biosynthesis |
| dre00061 | Fatty acid biosynthesis |
| dre00140 | Steroid hormone biosynthesis |
| dre00062 | Fatty acid elongation |
| dre00120 | Primary bile acid biosynthesis |
| dre01040 | Biosynthesis of unsaturated fatty acids |
| dre00030 | Pentose phosphate pathway |

**Figure S2:**
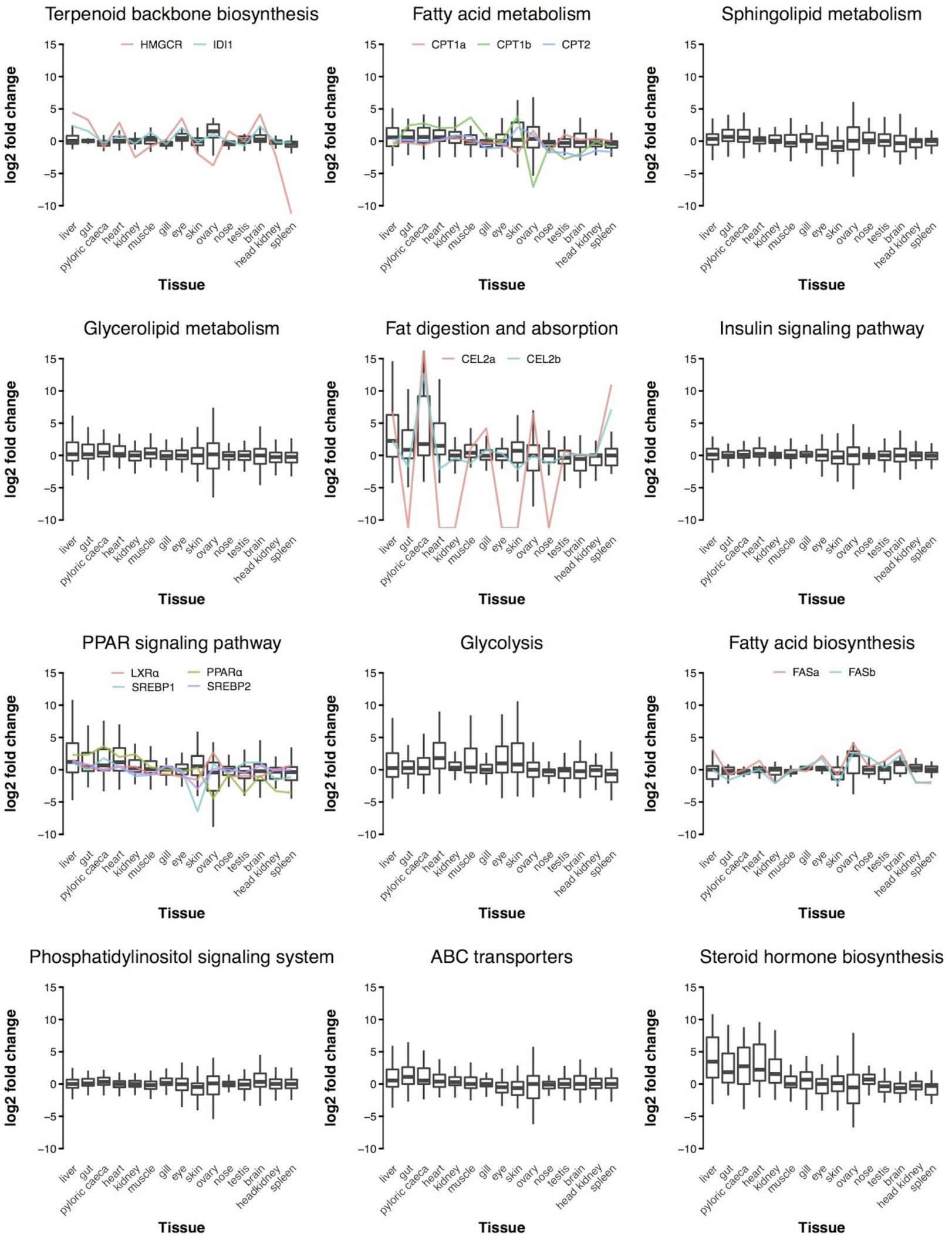
Tissue expression profiles of salmon genes in lipid metabolism pathways. Tissue expression profiles of salmon genes in KEGG pathways for *Salmo salar* tissues; pyloric caeca, liver, heart, gut, skin, kidney, muscle, eye, ovary, brain, nose, gill, testis, spleen, and head kidney. Gene expression levels are shown as the log2 fold change difference between each tissue FPKM to the medium FPKM across all tissues. Expression of select genes of interest are overlaid. These are 3-hydroxy-3-methyl-glutaryl-CoA reductase (HMGCR), isopentenyl-diphosphase Δisomerase (IDI1), carnitine palmitoyltransferase 1 (CPT1a and CPT1b) and 2 (CPT2), carboxyl ester lipase, tandem duplicate 2 (CEL2a and CEL2b), liver x receptor alpha (LXR*a*a), peroxisome proliferator-activated receptor alpha (PPARɑa), sterol regulatory element binding protein 1 (SREBP1) and 2 (SREBP2), and fatty acid synthase (FASa and FASb).

**Figure S3:**
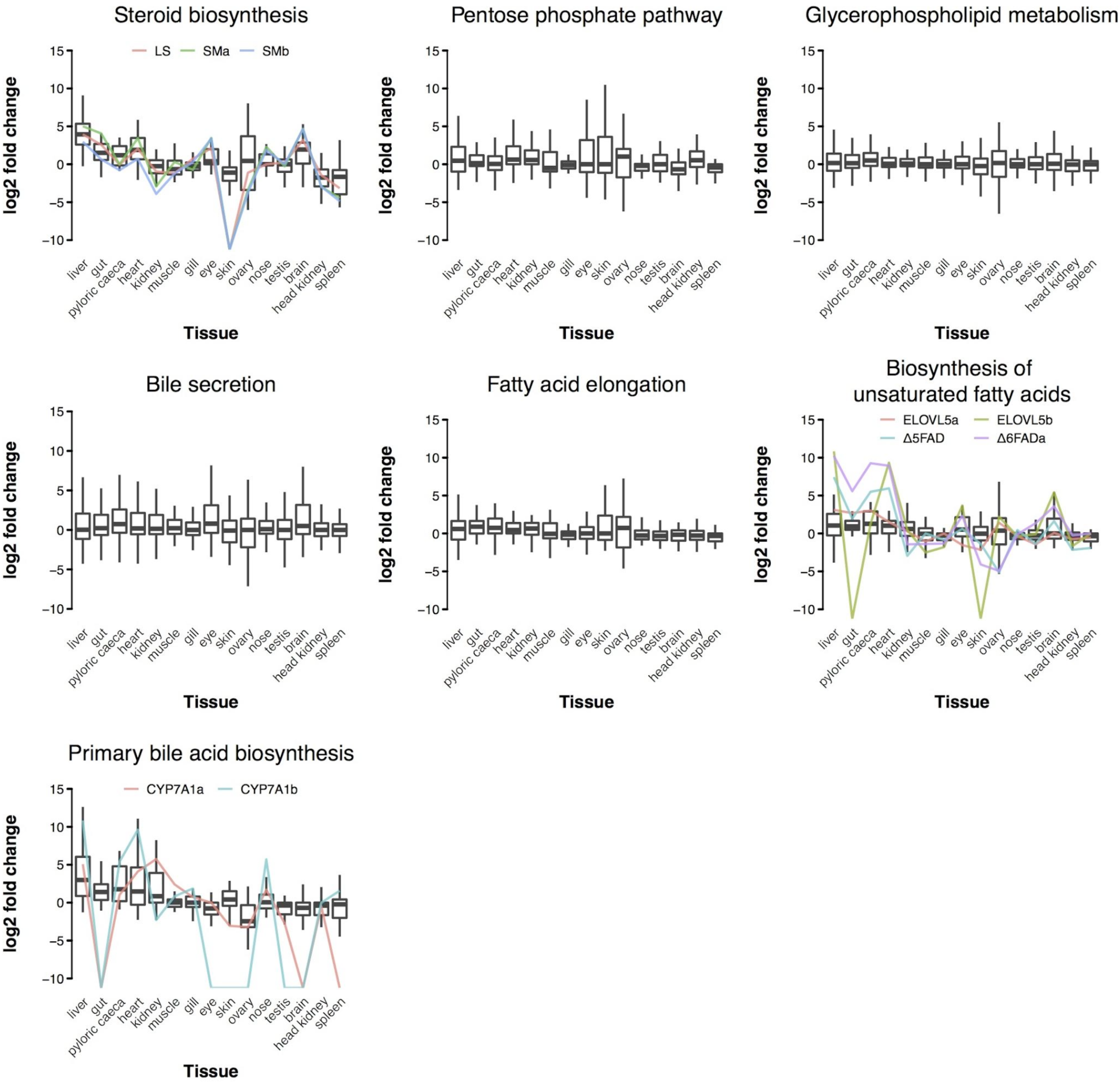
Tissue expression profiles of salmon genes in lipid metabolism pathways. Tissue expression profiles of salmon genes in KEGG pathways for *Salmo salar* tissues; pyloric caeca, liver, heart, gut, skin, kidney, muscle, eye, ovary, brain, nose, gill, testis, spleen, and head kidney. Gene expression levels are shown as the log2 fold change difference between each tissue FPKM to the medium FPKM across all tissues. Expression of select genes of interest are overlaid. These are lanosterol synthase (LS), squalene epoxidase (SMa and SMb), fatty acid elongase 5 (ELOVL5a and ELOVL5b), delta 5 fatty acid desaturase (Δ5FAD), delta 6 fatty acid desaturase (Δ6FAD), and cytochrome P450 7A1 (CYP7A1a and CYP7A1b).

**Table S3:**
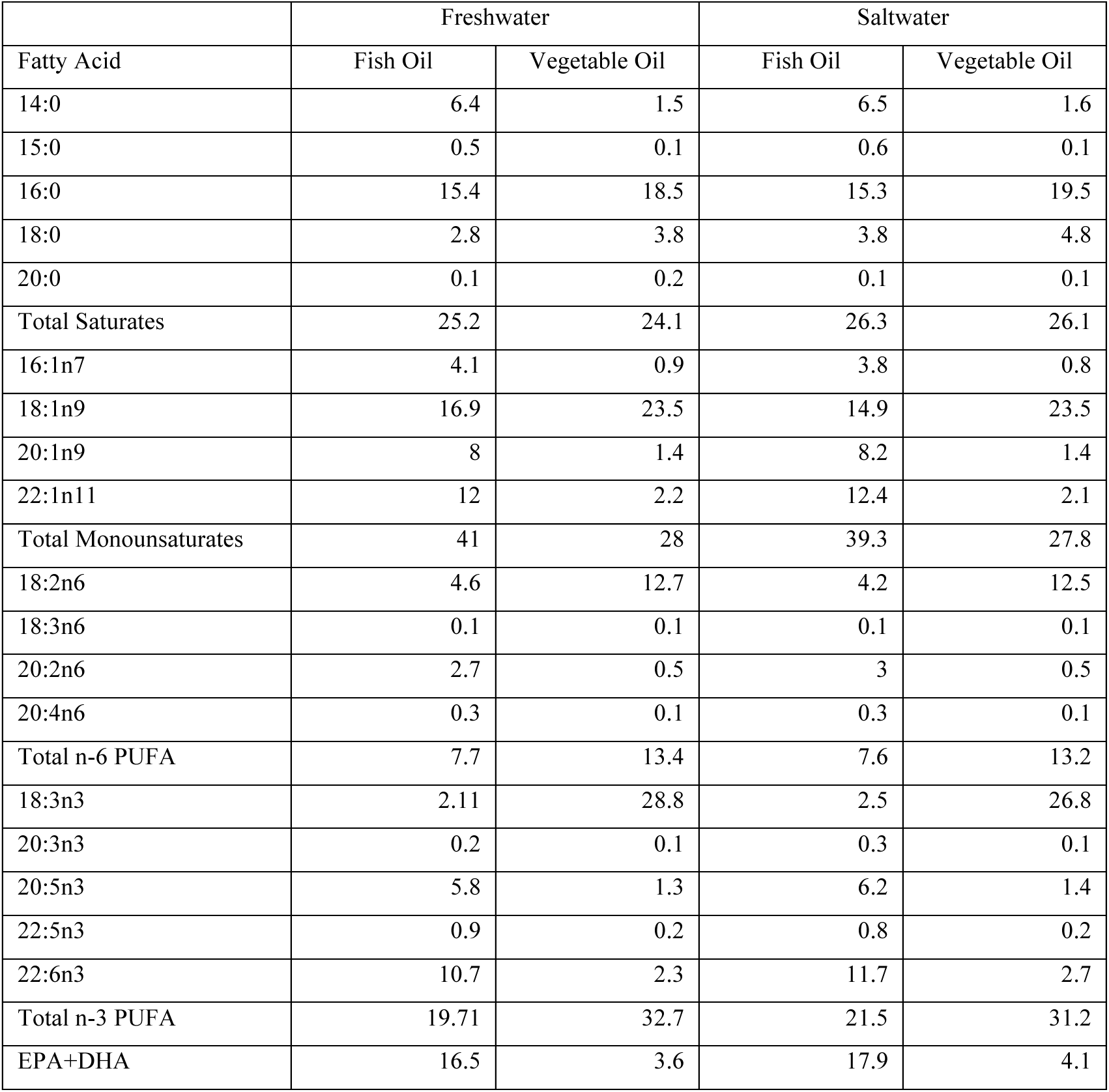
Fatty acid composition of feeds. All values are expressed as mass percentage of lipid fraction.

**Table S4:** Counts and proportions (%) of differentially expressed genes (DEG) for each lipid metabolism pathway. Results given for freshwater (FW) and saltwater (SW) DEGs from fish oil and vegetable oil feed comparison.

| Pathway name | Total genes<br>in pathway | FW<br>DEGs | SW<br>DEGs | FW&SW<br>DEGs | FW %<br>DEGs | SW %<br>DEGs | FW&SW<br>% DEGs |
| --- | --- | --- | --- | --- | --- | --- | --- |
| Phosphatidylinositol signaling system | 223 | 2 | 2 | 1 | 0.0090 | 0.0090 | 0.0045 |
| ABC transporters | 69 | 1 | 0 | 0 | 0.0145 | 0 | 0 |
| Sphingolipid metabolism | 105 | 1 | 1 | 0 | 0.0095 | 0.0095 | 0 |
| Glycolysis | 124 | 2 | 1 | 0 | 0.0161 | 0.0081 | 0 |
| Insulin signaling pathway | 304 | 5 | 3 | 0 | 0.0164 | 0.0099 | 0 |
| Bile secretion | 150 | 4 | 3 | 1 | 0.0267 | 0.0200 | 0.0067 |
| Glycerophospholipid metabolism | 183 | 3 | 5 | 0 | 0.0164 | 0.0273 | 0 |
| Fatty acid elongation | 49 | 2 | 1 | 0 | 0.0408 | 0.0204 | 0 |
| Primary bile acid biosynthesis | 28 | 1 | 1 | 0 | 0.0357 | 0.0357 | 0 |
| Glycerolipid metabolism | 108 | 4 | 4 | 0 | 0.0370 | 0.0370 | 0 |
| Steroid hormone biosynthesis | 59 | 3 | 4 | 1 | 0.0508 | 0.0678 | 0.0169 |
| PPAR signaling pathway | 114 | 9 | 8 | 5 | 0.0789 | 0.0702 | 0.0439 |
| Fat digestion and absorption | 66 | 2 | 5 | 0 | 0.0303 | 0.0758 | 0 |
| Pentose phosphate pathway | 56 | 5 | 1 | 0 | 0.0893 | 0.0179 | 0 |
| Biosynthesis of unsaturated fatty acids | 42 | 5 | 3 | 2 | 0.1190 | 0.0714 | 0.0476 |
| Fatty acid metabolism | 82 | 10 | 6 | 3 | 0.1220 | 0.0732 | 0.0366 |
| Fatty acid biosynthesis | 25 | 4 | 3 | 1 | 0.1600 | 0.1200 | 0.0400 |
| Terpenoid backbone biosynthesis | 34 | 11 | 0 | 0 | 0.3235 | 0 | 0 |
| Steroid biosynthesis | 30 | 17 | 5 | 3 | 0.5667 | 0.1667 | 0.1000 |

**Figure S4:**
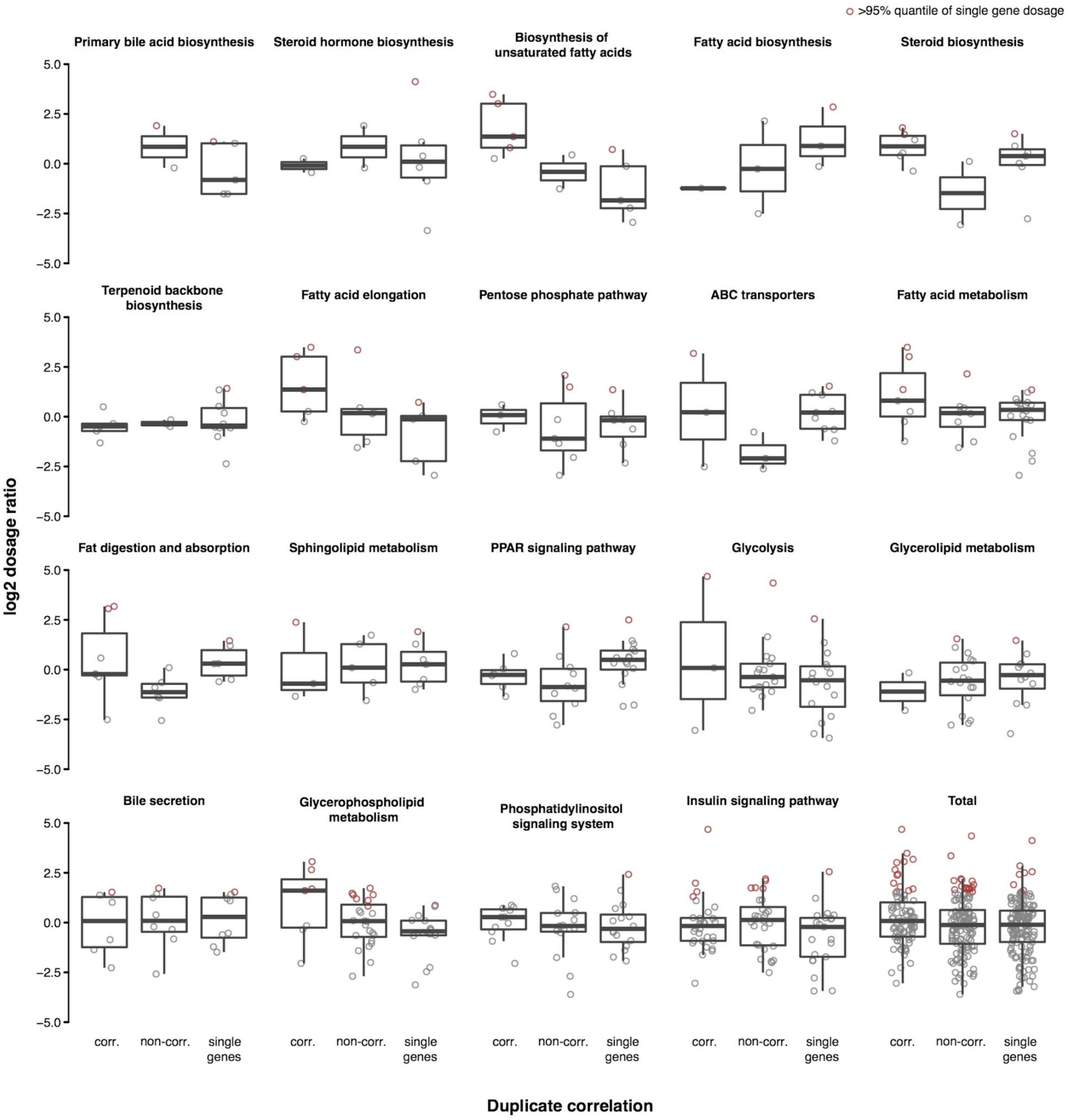
Gene dosage ratios and duplicate correlation for saltwater conditions. Log2 of gene dosage ratios of saltwater salmon ortholog expression in liver (FPKM, summed for duplicates) to expression of a pike ortholog, for total lipid metabolism genes and genes 19 KEGG pathways. Duplicates were grouped into correlated (corr.) or non-correlated (non-corr.) based on saltwater correlation results. Dosage ratios (points) greater than the 95% quantile of single gene dosages in the plot are marked in red.

**Figure S5:**
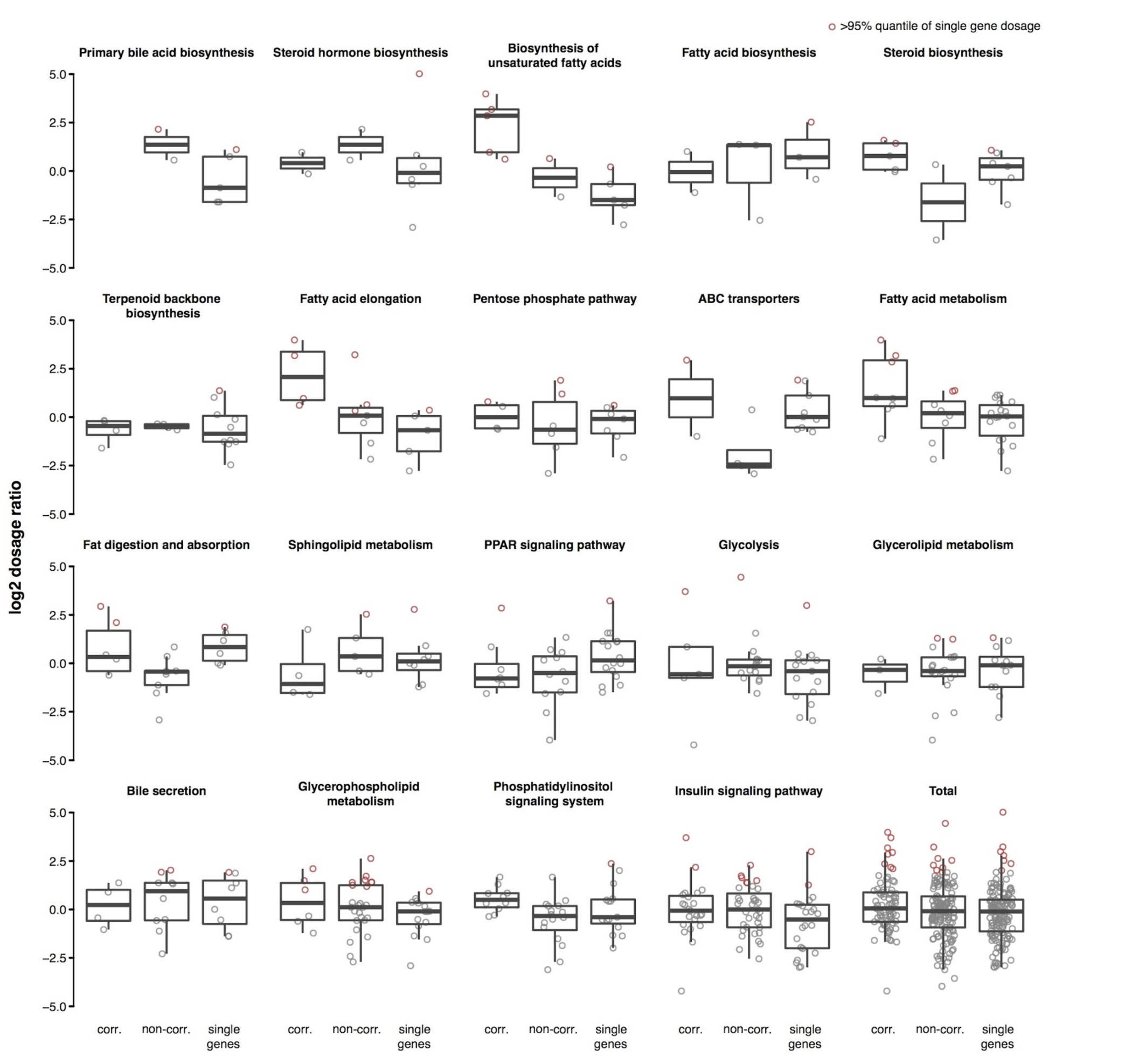
Gene dosage ratios and duplicate correlation for freshwater conditions. Log2 of gene dosage ratios of freshwater salmon ortholog expression in liver (FPKM, summed for duplicates) to expression of a pike ortholog, for total lipid metabolism genes and genes 19 KEGG pathways. Duplicates were grouped into correlated (corr.) or non-correlated (non-corr.) based on freshwater correlation results. Dosage ratios (points) greater than the 95% quantile of single gene dosages in the plot are marked in red.

